# The role of metal binding in the function of the human salivary antimicrobial peptide histatin-5

**DOI:** 10.1101/2022.01.07.472205

**Authors:** Louisa Stewart, YoungJin Hong, Isabel Holmes, Samantha Firth, Jack Bolton, Yazmin Santos, Steven Cobb, Nicholas Jakubovics, Karrera Djoko

## Abstract

Antimicrobial peptides (AMPs) are key components of diverse host innate immune systems. The family of human salivary AMPs known as histatins bind Zn and Cu. Fluctuations in Zn and Cu availability play significant roles in the host innate immune response (so-called “nutritional immunity”). Thus, we hypothesised that histatins contribute to nutritional immunity by influencing host Zn and/or Cu availability. We posited that histatins limit Zn availability (promote bacterial Zn starvation) and/or raise Cu availability (promote bacterial Cu poisoning). To test this hypothesis, we examined the interactions between histatin-5 (Hst5) and Group A *Streptococcus* (GAS), which colonises the human oropharynx. Our results showed that Hst5 does not strongly influence Zn availability. Hst5 did not induce expression of Zn-responsive genes in GAS, nor did it suppress growth of mutant strains that are impaired in Zn transport. Biochemical examination of purified peptides confirmed that Hst5 binds Zn only weakly. By contrast, Hst5 bound Cu tightly and it strongly influenced Cu availability. However, Hst5 did not promote Cu toxicity. Instead, Hst5 suppressed expression of Cu-inducible genes, stopped intracellular accumulation of Cu, and rescued growth of a Δ*copA* mutant strain that is impaired in Cu efflux. We thus proposed a new role for salivary histatins as major Cu buffers in saliva that contribute to microbial homeostasis in the oral cavity and oropharynx by reducing the potential negative effects of Cu exposure (*e*.*g*. from food) to microbes. Our results raise broad questions regarding the physiological roles of diverse metal-binding AMPs and the management of host metal availability during host-microbe interactions.

## INTRODUCTION

Antimicrobial peptides (AMPs) are short, often cationic peptides that are secreted by diverse organisms from across the domains of life^1^. These peptides usually act as immune effectors that kill invading microbes as part of the host innate immune system, but many also play key functions in the normal biology of the host organism. A sub-family of AMPs binds metals. These metallo-AMPs often synergise with metal ions or become activated upon metal binding^2-4^.

Salivary histatins comprise a family of His-rich, metallo-AMPs that are all derived from two parent peptides, namely Histatin-1 and Histatin-3^5,6^. Both histatins are expressed constitutively by the salivary glands of humans and some higher primates^6,7^. Upon secretion into the oral cavity, histatins are rapidly processed into shorter fragments^8,9^ by unidentified human salivary proteases or proteases from resident oral microbes. Of these fragments, Histatin-5 (Hst5; Table 1) is the best characterised.

**Table 1.**
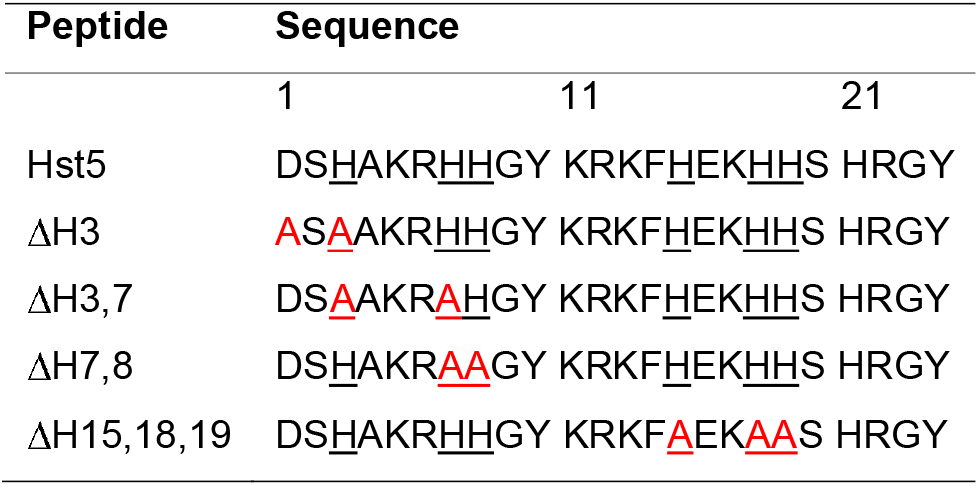
Hst5 peptides and variants used in this work.

Hst5 is noted for its direct antimicrobial activity against the fungus *Candida albicans*^10,11^. Unlike other AMPs, Hst5 does not appear to permeabilise fungal membranes, although it does destabilise some bacterial membranes^11^. Beyond its direct action on membranes, the antimicrobial activity of Hst5 requires the peptide to be internalised into the cytoplasm, usually *via* energy-dependent pathways for peptide uptake^11,12^. Once in the cytoplasm, Hst5 encounters its targets and causes toxicity *via* multiple pathways that are not fully elucidated^10,13^.

Hst5 contains the characteristic Zn-binding motif His-Glu-x-His-His (Table 1), but whether Zn binding is essential for the antimicrobial activity of this AMP is unclear. Hst5 derivatives that lack one or all three His residues remain active against *C. albicans*^14^. Conflicting reports show that addition of Zn can both enhance^15^ and suppress^16^ Hst5 activity against this fungus. The reason for this discrepancy has not been identified. In addition, Hst5 possesses three Cu-binding motifs, namely the N-terminal ATCUN motif that binds Cu(II) and two *bis*-His motifs that bind one Cu(I) each (Table 1)^17^. Addition of Cu potentiates the activity of Hst5 against *C. albicans*^17^. This potentiation relies on the Cu(I) site but not the Cu(II) site^17^.

Beyond histatins and metallo-AMPs, metal-dependent host innate immune responses are well described. In response to microbial infection, metal levels and those of metal-binding or metal-transport proteins within a host organism can rise and fall, leading to fluctuations in metal availability within different niches in the infected host. Increases in metal availability promote microbial poisoning while decreases in metal availability promote microbial starvation. These antagonistic host responses are known as “nutritional immunity”^18^. Do histatins and other metallo-AMPs contribute to these metal-dependent immune responses and, if so, how?

This study explored the relationship between Hst5 and metals, particularly Zn and Cu, and examined the role of this AMP in influencing metal availability during nutritional immunity. Based on the reported metal-dependent effects of Hst5 against *C. albicans* and on established features of nutritional immunity, we hypothesised that Hst5 either limits Zn availability (and promotes microbial Zn starvation) and/or raises Cu availability (and promotes Cu poisoning).

To test our hypothesis, the Gram-positive bacterium *Streptococcus pyogenes* (Group A *Streptococcus*, GAS) was used as a model. GAS colonises the human oropharynx, where it comes into contact with saliva and salivary components, but its interactions with Hst5 have not been described previously. Moreover, pathways for metal homeostasis in GAS are relatively well understood and phenotypes of mutant strains lacking key metal transport proteins are known^19^. These features enabled GAS to be exploited here as a tractable, well-defined experimental tool for examining the metal-dependent effects of Hst5.

## RESULTS

### Hst5 does not exert direct antibacterial effects against GAS or other oral streptococci

The effects of Hst5 on growth of GAS were examined in a metal-deplete, chemically defined medium (CDM)^20^. In this medium, up to 50 µM Hst5 (*ca*. total histatin concentrations in fresh salivary secretions^9^) did not affect growth of wild-type GAS (Figure 1A). Identical results were obtained in THY medium (Figure S1A).

**Figure 1.**
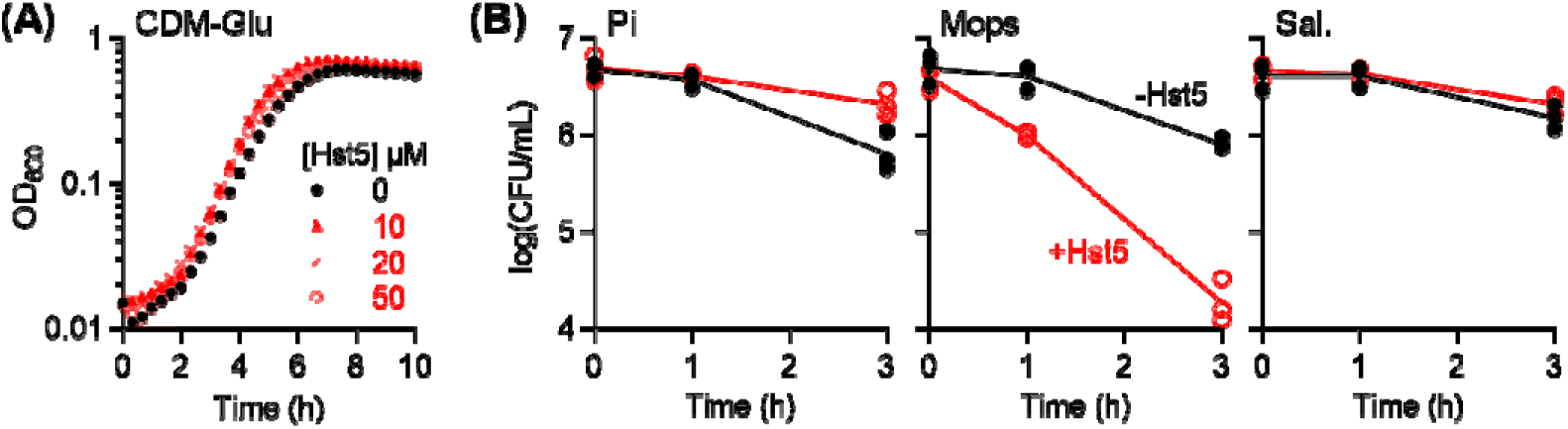
Effects of Hst5 on (A) growth and (B) survival of GAS. **(A)** Bacteria (*N =* 2) were cultured in CDM in the presence of Hst5 (0–50 µM). **(B)** Bacteria (*N =* 3) were incubated in phosphate buffer (10 mM, pH 7.4; Pi), Na-Mops buffer (10 mM, pH 7.4; Mops), or artificial saliva salts (pH 7.2-7.4; Sal., see Dataset S1a for composition), with (○) or without (●) Hst5 (50 µM).

The effects of Hst5 on GAS survival were examined in 10 mM phosphate buffer^11,15^. Under these conditions, up to 50 µM Hst5 did not kill GAS (Figure 1B). Instead, Hst5 prolonged survival of this bacterium (Figure 1B). A parallel control experiment showed that the same concentrations of Hst5 killed *Pseudomonas aeruginosa* within minutes^11^ (Figure S2A), confirming that our peptide stocks were active.

Like other cationic AMPs, the antimicrobial activity of Hst5 relies on initial electrostatic binding of the peptide to microbial surface proteins or membranes^11,21^. Such interactions are suppressed by salts and high ionic strength buffers^11,14,22-26^. To lower the ionic strength in our experiments, phosphate was replaced with Mops. Under these new conditions, Hst5 *did* kill GAS (Figure 1B). However, carryover salts from solutions used in preparing the inoculum abolished this killing effect (Figure S2B), underscoring the sensitivity of these assays to salt.

To better reflect the physiological context in which Hst5 plays a role, we repeated the kill assay in buffered “artificial saliva salts”, whose salt composition approximates healthy saliva (Dataset S1a). Hst5 did *not* kill GAS under these conditions (Figure 1B), confirming that the results in phosphate buffer are more physiologically relevant. For ease of comparison with existing literature, further experiments described below used phosphate buffer.

The lack of a direct antibacterial effect against GAS adds to the list of contradictory effects of Hst5 against streptococci reported in the literature^27-32^. Given the sensitivity of these assays to the specific experimental conditions, the effects of Hst5 on GAS and five oral streptococci, namely *S. anginosus, S. mutans, S. oralis, S. salivarius*, and *S. sanguinis*, were examined here in parallel. Hst5 did *not* kill or inhibit growth of any of the streptococci under these conditions (Figure 2, Figure S1B). In fact, Hst5 promoted growth of *S. anginosus, S. oralis*, and, to a lesser extent, *S. sanguinis*. The mechanism behind this growth-promoting activity of Hst5 is beyond the scope of the present work, but is presumably related to the metal-chelating ability of this AMP (described below).

**Figure 2.**
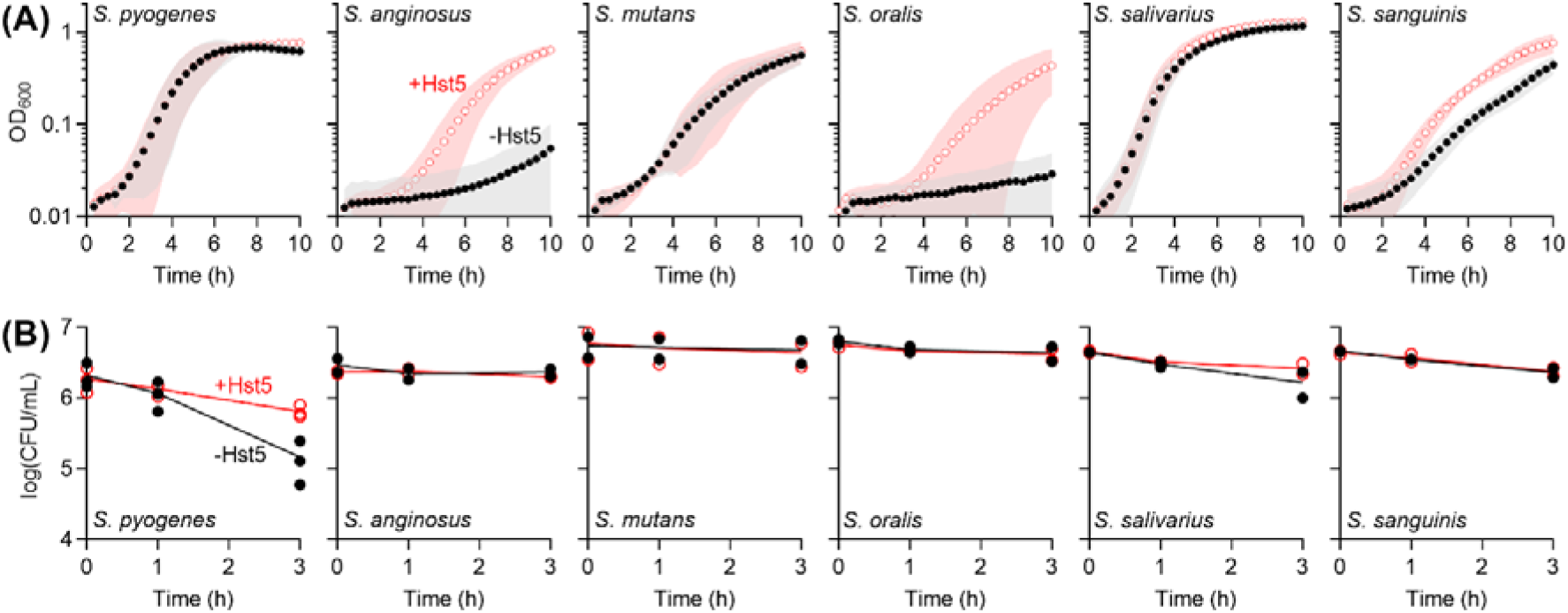
Effects of Hst5 on (A) growth and (B) survival of oral streptococci. Bacteria (*N =* 2) were cultured in CDM **(A)** or incubated in phosphate buffer **(B)**, with (○) or without (●) Hst5 (50 µM).

### Hst5 does not strongly influence Zn availability

To determine whether Hst5 contributes to nutritional immunity, the effects of this AMP on GAS were re-examined in the presence of up to 50 µM Zn (equimolar with Hst5 and in excess of Zn concentrations in whole saliva^33^). Zn neither suppressed nor enhanced the direct effects of Hst5 on GAS (Figure 3, Figure S1C), suggesting that Hst5 promotes neither Zn starvation nor poisoning, respectively.

**Figure 3.**
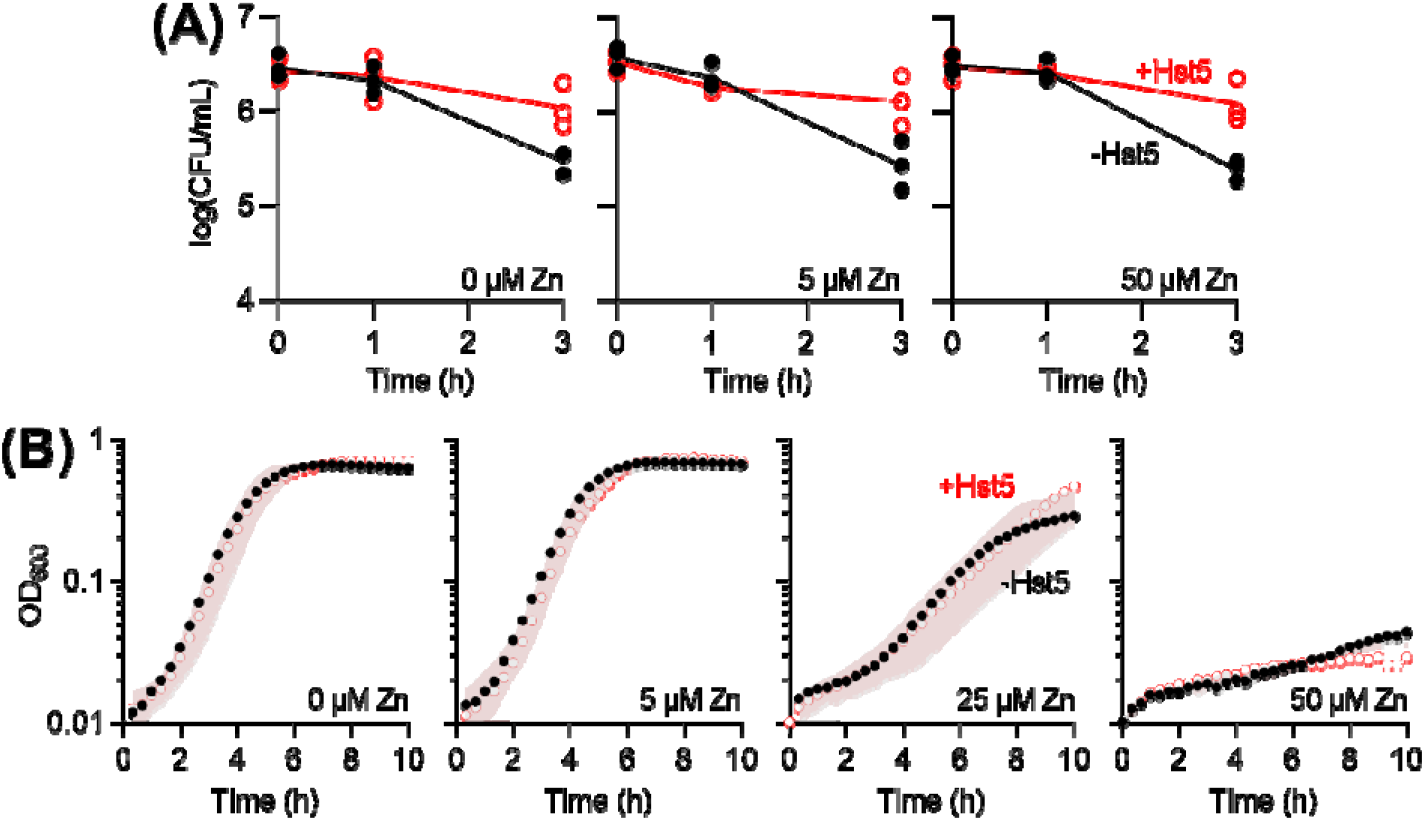
Effects of Zn on (A) survival and (B) growth of GAS in the presence of Hst5. Bacteria (*N =* 3) were incubated in phosphate buffer **(A)** or cultured in CDM **(B)**, in the presence of Zn (0–50 µM), with (○) or without (●) Hst5 (50 µM).

Zn starvation or poisoning may not strongly affect wild-type GAS since the transcriptionally-responsive system for Zn homeostasis responds to, and thus counters, such perturbations in Zn availability (Figure 4). This transcriptional response, *i*.*e*. expression of the three Zn-responsive genes *adcAI, adcAII*, and *czcD* (Figure 4), was examined here. Only bacteria grown in CDM were used for analyses, since poor RNA yields were obtained from bacteria that were incubated in phosphate buffer, likely associated with the progressive loss of viability under these conditions (*cf*. Figure 1B).

**Figure 4.**
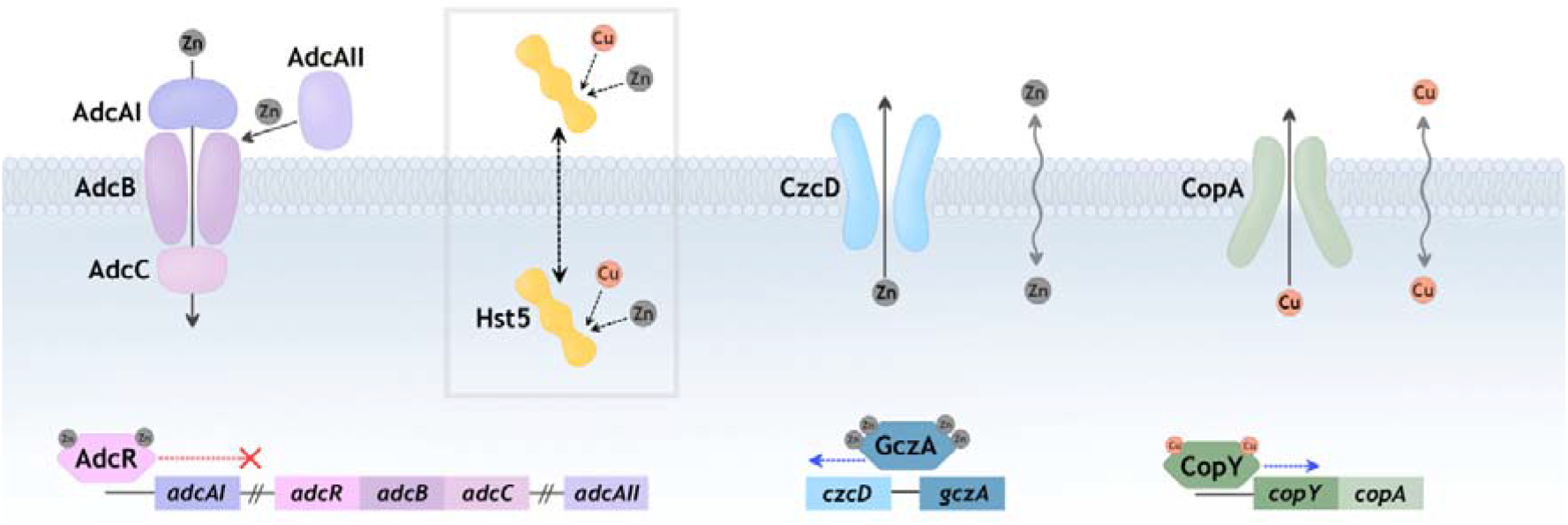
Metal homeostasis in GAS and hypothesised actions of Hst5. **Zn uptake:** AdcAI and AdcAII capture extracellular Zn and transfer this metal to AdcBC for import into the cytoplasm. These proteins are transcriptionally upregulated by AdcR in response to decreases in Zn availability^34^. **Zn efflux:** CzcD exports excess Zn out of the cytoplasm. It is transcriptionally upregulated by GczA in response to increases in Zn availability^35^. **Cu efflux:** CopA exports excess Cu out of the cytoplasm. It is transcriptionally upregulated by CopY in response to increases in Cu availability^36^. **Hypothesised actions of Hst5:** Hst5 may remain extracellular, bind Zn or Cu, and suppress extracellular metal availability. Alternatively, Hst5 may become internalised and suppress intracellular metal availability. Hst5 may also become internalised as the Zn-Hst5 or Cu-Hst5 complex, facilitate entry of Zn or Cu into the cytoplasm, and increase metal availability.

In the control experiment, adding Zn alone did not further repress transcription of *adcAI* and *adcAII*, but it did induce expression of *czcD* (Figure S3A), consistent with an increase in Zn availability. Conversely, adding the Zn chelator TPEN did not affect transcription of *czcD*, but it did induce expression of *adcAI* and *adcAII*, consistent with a decrease in Zn availability (Figure S3B). By contrast, adding Hst5 alone perturbed neither the basal expression of *adcAI* or *adcAII* (Figure 5A), nor the Zn-dependent expression of *czcD* (Figure 5B).

**Figure 5.**
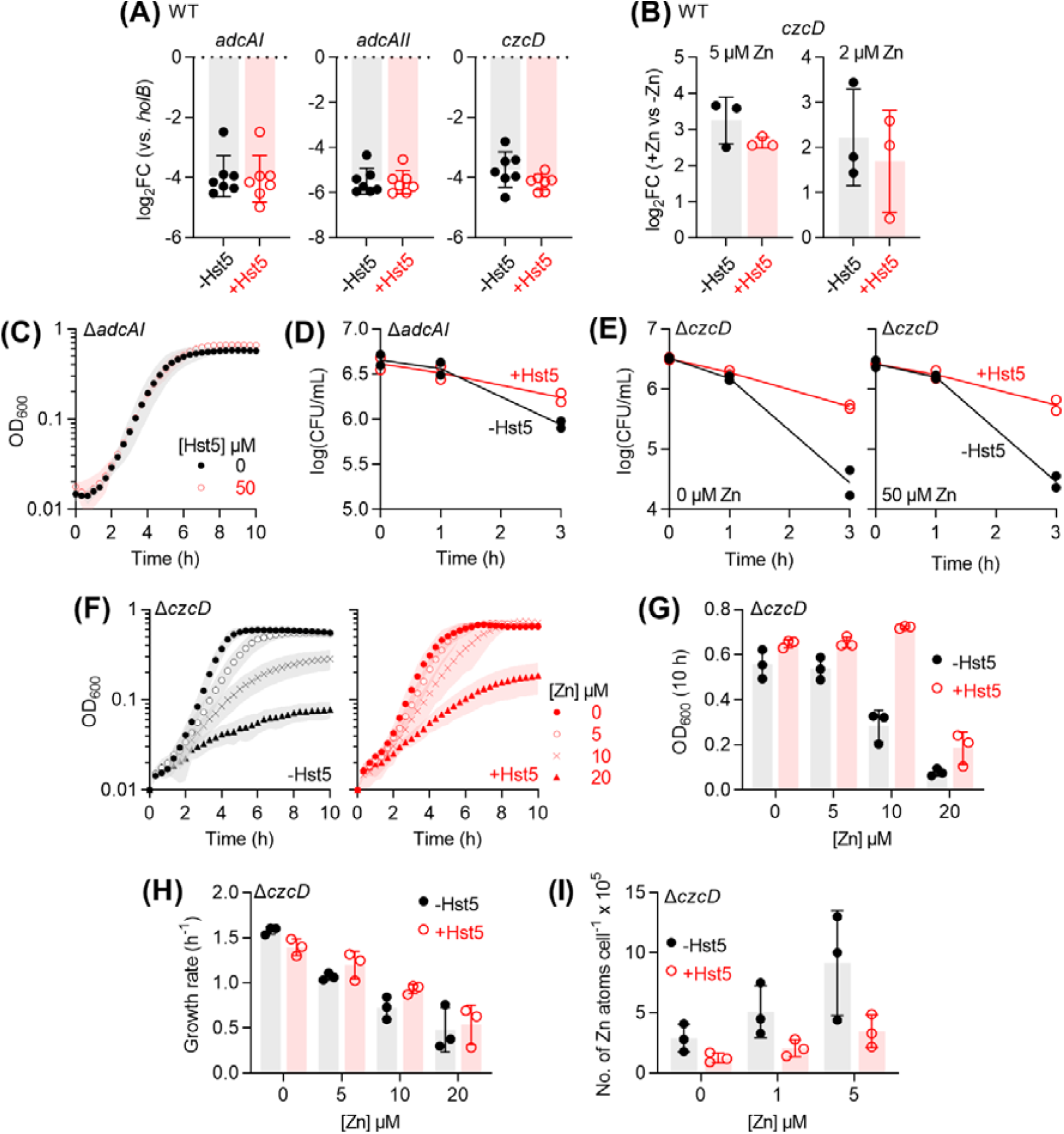
Effects of Hst5 on Zn availability. **(A) Expression of Zn-responsive genes**. Bacteria (*N =* 7) were cultured in CDM with (○) or without (●) Hst5 (50 µM). Levels of *adcAI, adcAII*, and *czcD* mRNA were determined by qRT-PCR and normalised to *holB*. **(B) Zn-dependent expression of *czcD***. Bacteria (*N =* 3) were cultured in CDM with or without added Zn (2 or 5 µM), with (○) or without (●) Hst5 (50 µM). Levels of *czcD* mRNA were measured by qRT-PCR, normalised to *holB*, and compared to normalised mRNA levels of the corresponding untreated controls (0 µM added Zn). **(C) Growth of** Δ***adcAI***. Bacteria (*N =* 3) were cultured in CDM with or without Hst5 (0 or 50 µM). **(D) Survival of** Δ***adcAI***. Bacteria (*N =* 2) were incubated in phosphate buffer with (○) or without (●) Hst5 (50 µM). **(E) Survival of** Δ***czcD***. Bacteria (*N =* 2) were incubated in phosphate buffer with or without added Zn (50 µM), with (○) or without (●) Hst5 (50 µM). **(F) Growth of** Δ***czcD***. Bacteria (*N =* 3) were cultured in CDM in the presence of Zn (0–20 µM), with (○) or without (●) Hst5 (50 µM). **(G) Final culture densities from panel F. (H) Exponential growth rates from panel F. (I) Intracellular Zn levels in** Δ***czcD***. Bacteria (*N =* 3) were cultured in CDM in the presence of Zn (0–5 µM), with or without Hst5 (50 µM). Intracellular levels of Zn were measured by ICP MS and normalised to colony counts.

As described earlier, Zn uptake by AdcAI and Zn efflux by CzcD may mask the effects of Hst5 on Zn availability (Figure 4). Thus, the effects of Hst5 were examined further using the Δ*adcAI* and Δ*czcD* mutant strains. Although additional Zn-binding lipoproteins such as AdcAII contribute to Zn acquisition^37^, AdcAI is thought to act as the primary Zn importer^37,38^. Therefore, only the Δ*adcAI* mutant was employed here. The control experiment confirmed that the Δ*adcAI* and Δ*czcD* mutant strains were sensitive to growth inhibition by TPEN^37,38^ and added Zn^35,38^, respectively (Figure S4).

The Δ*adcAI* mutant strain displayed wild-type growth and survival phenotypes in the presence of Hst5 (Figures 5B–C), strengthening the proposal that Hst5 does not starve GAS of nutrient Zn.

Likewise, the Δ*czcD* mutant strain displayed wild-type survival phenotype (Figure 5E). Interestingly, Hst5 weakly improved (instead of further inhibited) growth of the Δ*czcD* mutant in the presence of 10 µM of added Zn (Figure 5F). This growth-promoting effect became apparent only upon comparing final culture densities (Figure 5G), since exponential growth rates remained unchanged (Figure 5H). It appeared to require the predicted Zn-binding ligands His15, His18, and His19^39,40^, since growth of the Zn-treated Δ*czcD* mutant in the presence of the ΔH15,18,19 variant of Hst5 was indistinguishable with growth in the absence of Hst5 (Figure S5A-B). The roles of the other His residues were less clear (Figure S5A-B). Nevertheless, it can be concluded that Hst5 suppresses (instead of potentiates) Zn toxicity to GAS.

Two mechanisms are immediately plausible (*cf*. Figure 4): (i) Hst5 binds extracellular Zn and weakly suppresses entry and accumulation of this metal ion into the cytoplasm, leading to less Zn toxicity, or (ii) Hst5 binds intracellular Zn and enables more Zn to accumulate in the cytoplasm, but with less toxicity. To distinguish these models, total intracellular Zn levels in the Δ*czcD* mutant strain were assessed by ICP MS. Only up to 5 µM Zn was used, since adding 10 µM Zn did not produce sufficient biomass for metal analyses. Only wild-type Hst5 was used, owing to the large culture volumes required and the high cost of peptide synthesis.

Figure 5I shows that growth in the presence of added Zn increased intracellular Zn levels in the Δ*czcD* mutant, but co-treatment with Hst5 suppressed this effect. These results initially appeared to support the first model, in which Hst5 binds extracellular Zn. However, intracellular Cu levels in these samples were similarly elevated in the absence of Hst5, and similarly suppressed in the presence of Hst5 (Figure S5C). At this stage, we cannot exclude the possibility that Zn treatment led to spurious effects associated with the observed growth defect. Thus, while our data hint at a role for Hst5 in weakly influencing extracellular Zn availability to GAS, they are not conclusive, particularly when compared with the clear role of Hst5 in strongly influencing Cu availability (described below).

### Hst5 binds extracellular Cu(II) and strongly limits Cu availability

Like Zn, adding up to 50 µM of Cu (equimolar with Hst5; *ca*. 10X higher than Cu concentrations in saliva^41-43^) did not directly affect the growth or survival phenotype of wild-type GAS in the presence of Hst5 (Figure 6, Figure S1D). However, as in the case with Zn, any effect of Hst5 on Cu availability may not directly affect wild-type GAS as a result of the transcriptionally-responsive system for Cu export (Figure 4). This transcriptional response was hereby examined to probe the Cu-linked action of Hst5.

**Figure 6.**
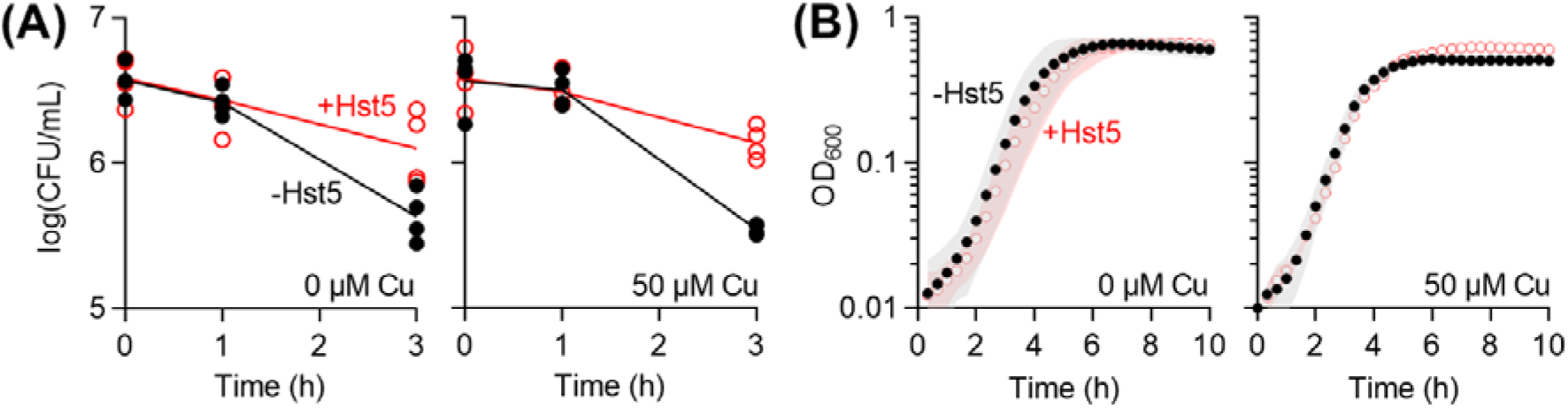
Effects of Cu on (A) survival and (B) growth of GAS in the presence of Hst5. **(A)** Bacteria (*N =* 5) were incubated in phosphate buffer **(A)** or cultured in CDM glucose **(B)**, with or without added Cu (50 µM), with (○) or without (●) Hst5 (50 µM).

The control experiment showed that adding the extracellular Cu chelator BCS did not further repress expression of *copA* and *copZ* (Figure S3C), suggesting that GAS grown in CDM was Cu-deplete. Adding Cu to the culture medium induced expression of both genes (Figure S3D, Figure 7A), consistent with an increase in Cu availability^20^. Intriguingly, co-treatment with Hst5 suppressed (instead of enhanced) this Cu-dependent induction (Figure 7A). This effect required the predicted Cu(II) binding site^17,44^, since the ΔH3 and ΔH3,7 variants lacking His3 (Table 1) were less effective at reducing expression of *copA* and *copZ* (Figure 7B). By contrast, it did not require the predicted Cu(I) binding sites^17^, since the effects of the ΔH7,8 and ΔH15,18,19 variants lacking either of the *bis*-His motifs (Table 1) were indistinguishable to that of the wild-type peptide (Figure 7B).

**Figure 7.**
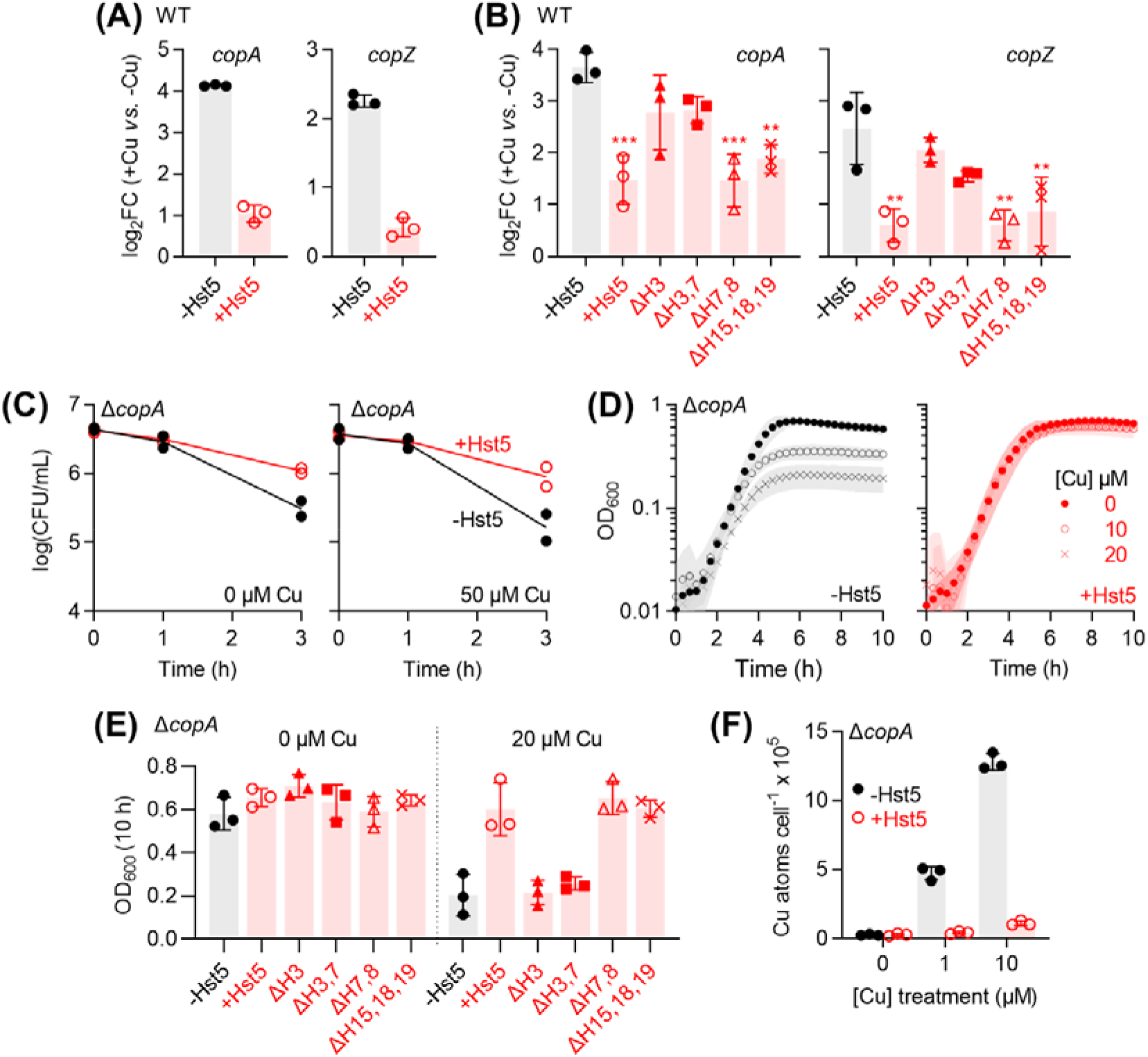
Effects of Hst5 on Cu availability. **(A) Expression of Cu-inducible genes**. Bacteria (*N =* 3) were cultured in CDM with or without added Cu (10 µM), with or without Hst5 (50 µM). Levels of *copA* and *copZ* mRNA were measured by qRT-PCR, normalised to *holB*, and compared with normalised mRNA levels of the corresponding untreated controls (0 µM Cu). **(B) Effects of Hs5 variants on expression of Cu-inducible genes**. The experiment was performed as described in panel A. The following treatments suppressed Cu-dependent copA expression when compared with untreated control: Hst5 (\*\*\**P* = 0.0003), ΔH7,8 (\*\*\**P* = 0.0003), ΔH15,18,19 (\*\**P* = 0.0018). The following treatments had no effect: ΔH3 (*P* = 0.1), ΔH3,7 (*P* = 0.2). The following treatments suppressed Cu-dependent copZ expression when compared with untreated control: Hst5 (\*\*\**P* = 0.001), ΔH7,8 (\*\*\**P* = 0.001), ΔH15,18,19 (\*\**P* = 0.003). The following treatments had no effect: ΔH3 (*P* = 0.7), ΔH3,7 (*P* = 0.1). **(C) Survival of** Δ***copA***. Bacteria (*N =* 2) were incubated in phosphate buffer with or without added Cu (50 µM), with (○) or without (●) Hst5 (50 µM). **(D) Growth of** Δ***copA***. Bacteria (*N =* 3) were cultured in CDM in the presence of added Cu (0–20 µM), with or without Hst5 (50 µM). **(E) Effects of Hst5 variants on growth of** Δ***copA***. Bacteria (*N =* 3) were cultured in CDM with or without added Cu (20 µM), with or without Hst5 or its variants (50 µM). Complete growth curves are shown in Figure S6. For ease of comparison, only OD_600_ values from the end of the experiment (*t* = 10 h) were plotted here. **(F) Intracellular Cu levels in** Δ***copA***. Bacteria (*N =* 3) were cultured in CDM in the presence of Cu (0–10 µM), with or without Hst5 (50 µM). Intracellular levels of Cu were measured by ICP MS and normalised to colony counts.

Further examination using a Cu-sensitive Δ*copA* mutant strain that lacks the Cu-effluxing P-type ATPase^20^ (Figure 4) revealed no difference between the survival phenotype of this mutant strain and that of the wild-type in the presence of Hst5 (Figure 7C). There was, however, a clear difference in their growth phenotypes. Co-treatment with Hst5 rescued growth of the Δ*copA* mutant strain in the presence of added Cu (Figure 7D). This protective effect again required the His3 ligand for Cu(II), but neither of the two *bis*-His ligands for Cu(I) (Figure 7E, Figure S6A). These results indicate that Hst5 acts as a Cu(II)-specific peptide.

Two mechanisms are again plausible (Figure 4): (i) Hst5 binds extracellular Cu and suppresses entry of Cu into the GAS cytoplasm, leading to less Cu toxicity, or (ii) Hst5 binds intracellular Cu, allowing intracellular Cu levels to rise without significant toxicity. The latter would resemble the model described for GSH in binding (buffering) excess intracellular Cu^20^. Since Cu(II) is not thought to exist within the reducing cytoplasm, the first model is more likely. Consistent with this proposal, ICP MS analyses of total metal levels in the Δ*copA* mutant confirmed that growth in the presence of Cu led to an increase in total intracellular Cu levels, but co-treatment with Hst5 strongly suppressed these levels (Figure 7F). Unlike the situation described earlier for the Δ*czcD* mutant, there were no unanticipated effects on other metal levels such as Zn (Figure S6B). Thus, it can be concluded that Hst5 binds extracellular Cu(II) and strongly limits (instead of promotes) Cu availability to GAS.

### Molecular basis for the action of Hst5 in weakly influencing Zn availability

Hst5 is thought to bind up to three Zn atoms. ITC measurements yielded log *K*_Zn_ values of 4.0, 5.0, and 5.1^44^. In agreement with Zn binding weakly to Hst5, the Zn-Hst5 complex dissociated upon passage through a desalting column (Figure 8A).

**Figure 8.**
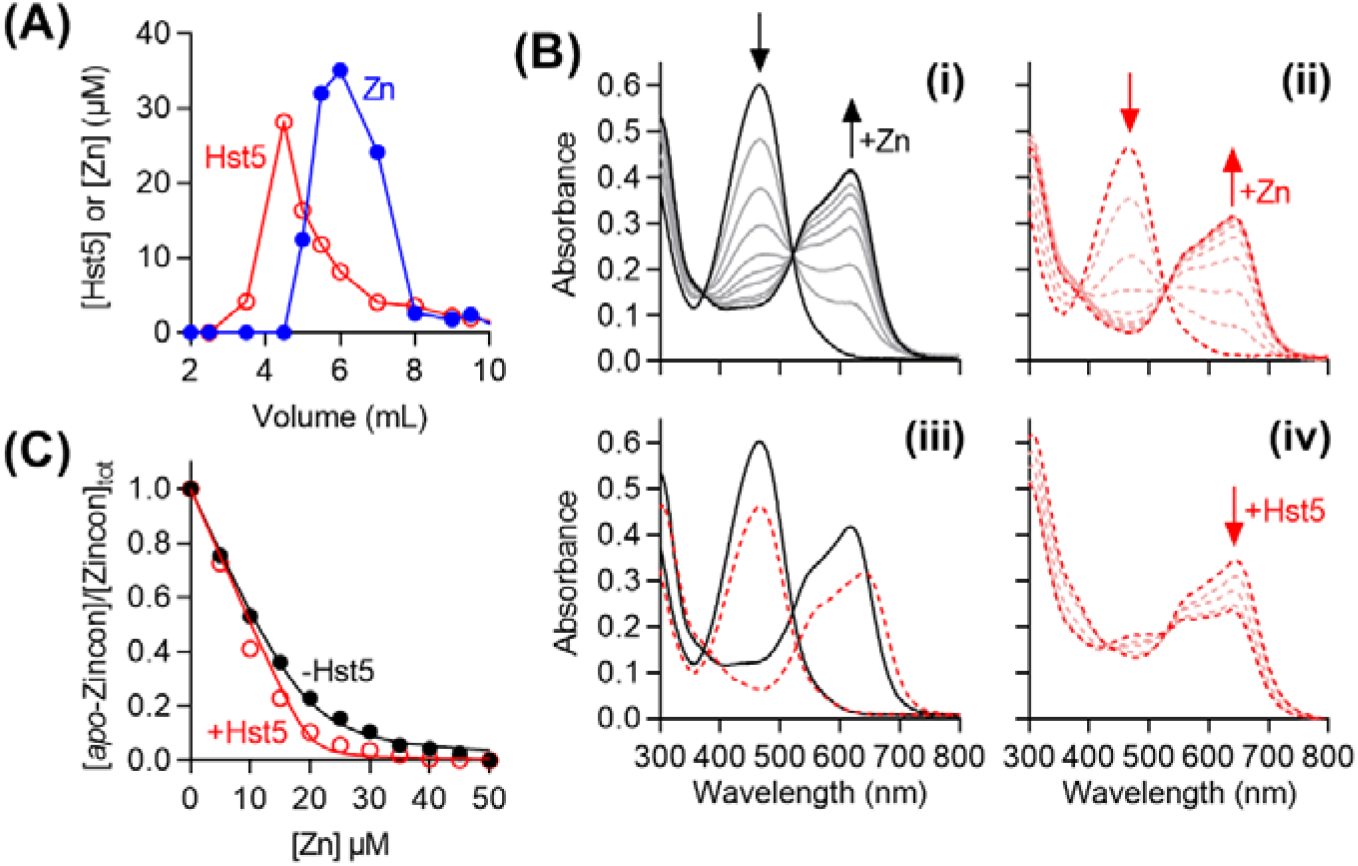
Zn affinity of Hst5. **(A)** Representative separation of Hst5 (○) and Zn (●) on a polyacrylamide desalting column. **(B)** Representative spectral changes upon addition of Zn (0–50 µM) into *apo*-Zincon (20 µM): **(i)** in the absence (solid traces) or **(ii)** presence (dashed traces) of Hst5 (20 µM). **(iii)** Overlaid spectra for 0 and 50 µM Zn from panels (i) and (ii). **(iv)** Representative spectral changes upon addition of excess Hst5 (0–200 µM) into a solution of Zn (20 µM) and *apo*-Zincon (25 µM). **(C)** Representative normalised plot of the absorbance intensities of *apo*-Zincon at 467 nm upon addition of Zn in the absence (●) or presence (○) of Hst5 (20 µM).

The affinities of Hst5 to Zn were re-examined here by equilibrium competition with the colorimetric Zn indicator Zincon (log *K*_Zn_ ∼ 6.0) and monitoring solution absorbances of *apo*-Zincon and Zn-Zincon at 466 nm and 620 nm, respectively (Figure 8B). Unexpectedly, the competition curve (in the presence of Hst5) was nearly indistinguishable from the control (in the absence of Hst5) (Figure 8C). Moreover, a new peak at 650 nm appeared in the presence of Hst5 (Figure 8B), indicating formation of a new species, likely a ternary complex between Hst5, Zincon, and Zn. This peak did not completely disappear upon adding excess Hst5 (Figure 8B). These results indicate that Hst5 does not compete effectively with Zincon, and that this peptide binds Zn more weakly than previously estimated by ITC^44^.

A previous study showed effective competition between Hst5 and Zincon in phosphate buffer, with Hst5 removing 2 molar equiv. of Zn from Zincon^15^. However, when used at millimolar concentrations, phosphate can compete for binding Zn (log *K*_Zn_ ∼ 2.4)^45^. Repeating the control titration in phosphate buffer (50 mM) instead of Mops led to clear partitioning of Zn between Zincon and the buffer (Figure S7A-B). Prolonged incubation (>10 min) of Zn-Zincon in this buffer led to loss of the characteristic blue colour (Figure S7C). For these reasons, estimates of Zn affinity and stoichiometry of Hst5 using Zincon in Mops buffer are likely to be more reliable.

The weak binding of extracellular Zn to Hst5 was clearly insufficient to starve wild-type GAS of nutrient Zn (*cf*. Figures 5A, 5C), indicating that this peptide does not compete with the high-affinity, Zn-specific uptake protein AdcAI (*cf*. Figure 4). Therefore, the Zn affinities of AdcAI were examined here by competition with the colorimetric Zn indicator Mag-fura2 (Mf2). The competition curve, generated by monitoring the solution absorbance of *apo*-Mf2 at 377 nm, clearly showed two Zn binding sites in AdcAI (Figure 9A) as anticipated^46^. The tight site outcompeted Mf2, as evidenced by the lack of Representative spectral changes upon adding up to 1 molar equiv. of Zn *vs*. AdcAI (Figure 9A). The weak site competed effectively with Mf2 with a log *K*_ZN =_ 8.5 (±0.2). The affinity of the tight site was better estimated using Quin-2 (Q2) as the competitor. By monitoring absorbance of *apo-*Q2 at 266 nm, a log *K*_ZN =_ 12.5 (±0.2) was obtained for this site (Figure 9B).

**Figure 9.**
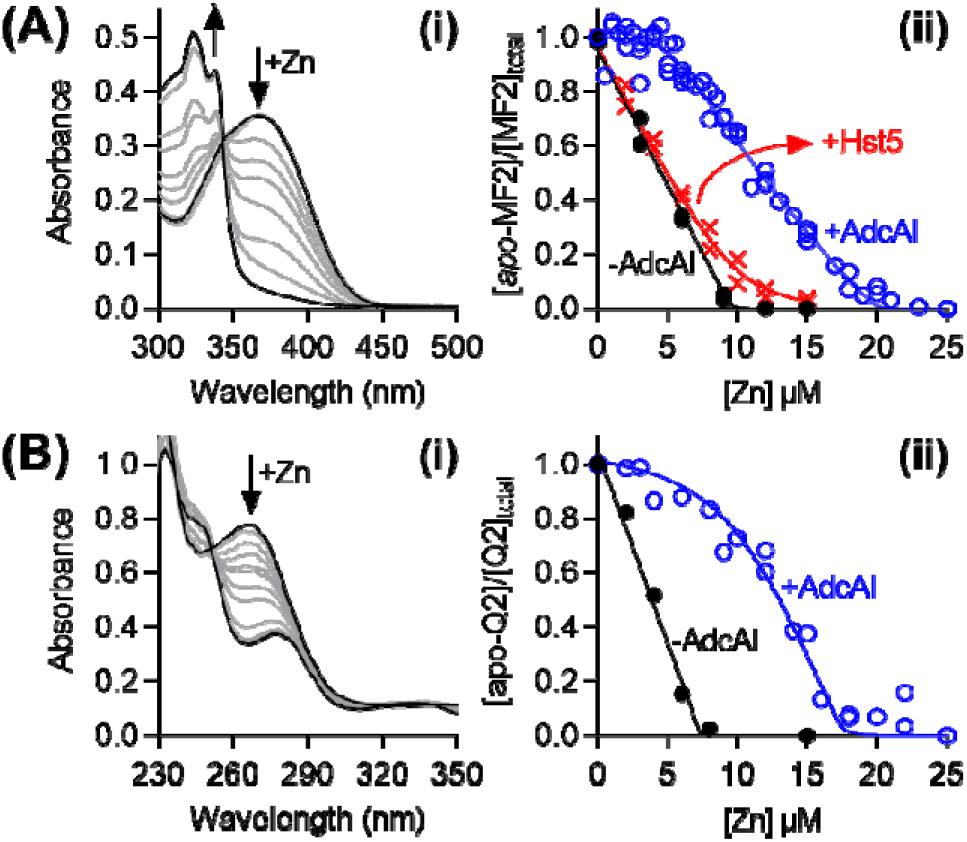
Zn affinity of AdcAI. **(A) Weak site. (i)** Representative spectral changes upon titration of Zn (0–25 µM) into a mixture of *apo*-Mf2 (10 µM) and AdcAI (5 µM). **(ii)** Normalised plot of the absorbance intensities of *apo*-MF2 (10 µM) at 377 nm upon addition of Zn in the absence (●) or presence (○) of AdcAI (5 µM). Competition with Hst5 (⍰; 10 µM) is shown for comparison. **(B) Tight site. (i)** Observed Representative spectral changes upon titration of Zn (0–25 µM) into a mixture of *apo*-Q2 (7.5 µM) and AdcAI (10 µM). **(ii)** Normalised plot of the absorbance intensities of *apo*-Q2 (7.5 µM) at 262 nm upon addition of Zn in the absence (●) or presence (○) of AdcAI (10 µM).

The log *K*_Zn_ values determined here were each *ca*. 1000-fold tighter than those determined previously by ITC^46^. ITC can underestimate metal binding affinities due to lack of sensitivity, lack of specificity, and potential side reactions (*e*.*g*. competition with buffers)^47^. Crucially, Hst5 did not compete with Mf2 for Zn (Figure 9A). These *relative* affinities, determined using the *same* approach under the *same* conditions, support the hypothesis that Hst5 does not compete with AdcAI for binding Zn, and provide a molecular explanation for why this AMP does not limit the availability of extracellular nutrient Zn to wild-type GAS.

Hst5 did not affect growth of GAS even when AdcAI was deleted by mutagenesis (*cf*. Figure 5D), suggesting that this peptide does not compete with other high-affinity Zn uptake proteins such as AdcAII (*cf*. Figure 4). AdcAII was expressed here for metal competition assays. However, consistent with a previous report^48^, it co-purified with 1 molar equiv. of bound Zn, which could not be removed without denaturing the protein. Nevertheless, the reported affinity of the *S. pneumoniae* homologue (log *K*_ZN =_ 7.7; 67% identity, 81% similarity), determined *via* competition with Mf2^49^, is consistent with our proposal that Hst5 does not compete effectively with AdcAII for binding Zn.

### Molecular basis for the action of Hst5 in strongly influencing Cu availability

Hst5 binds one Cu(II) ion with log *K*_Cu_ = 11.1, as determined previously by competition with NTA^17^. Since both Hst5 and NTA have weak optical signals (Figure S8), this log *K*_Cu_ value was re-evaluated here using PAR as an intensely coloured mediator^50^ between NTA and Hst5. The control titration confirmed that adding Cu(II) decreased the solution absorbance of *apo-*PAR at 400 nm and concomitantly increased that of the Cu(II)-PAR complex^51^ at 512 nm (Figure 10A, Figure S9A). PAR was then competed with 20 molar equiv. of NTA and, separately, Hst5 (Figures 10B-C). This PAR-mediated competition with NTA yielded a log *K*_Cu_ = 12.1 (±0.1) for Hst5 (Figure 10C, Figure S9B-C), *i*.*e*. ∼10-fold tighter than estimates from the direct competition with NTA^17^. As previously acknowledged, the weak solution absorbances of Cu^II^-NTA and Cu^II^-Hst5 complexes did not saturate even in the presence of excess Cu (Figure S8B), indicating potential Cu-buffer interactions that were not accounted in the calculations, which may explain the slight underestimate in the literature^17,52^.

**Figure 10.**
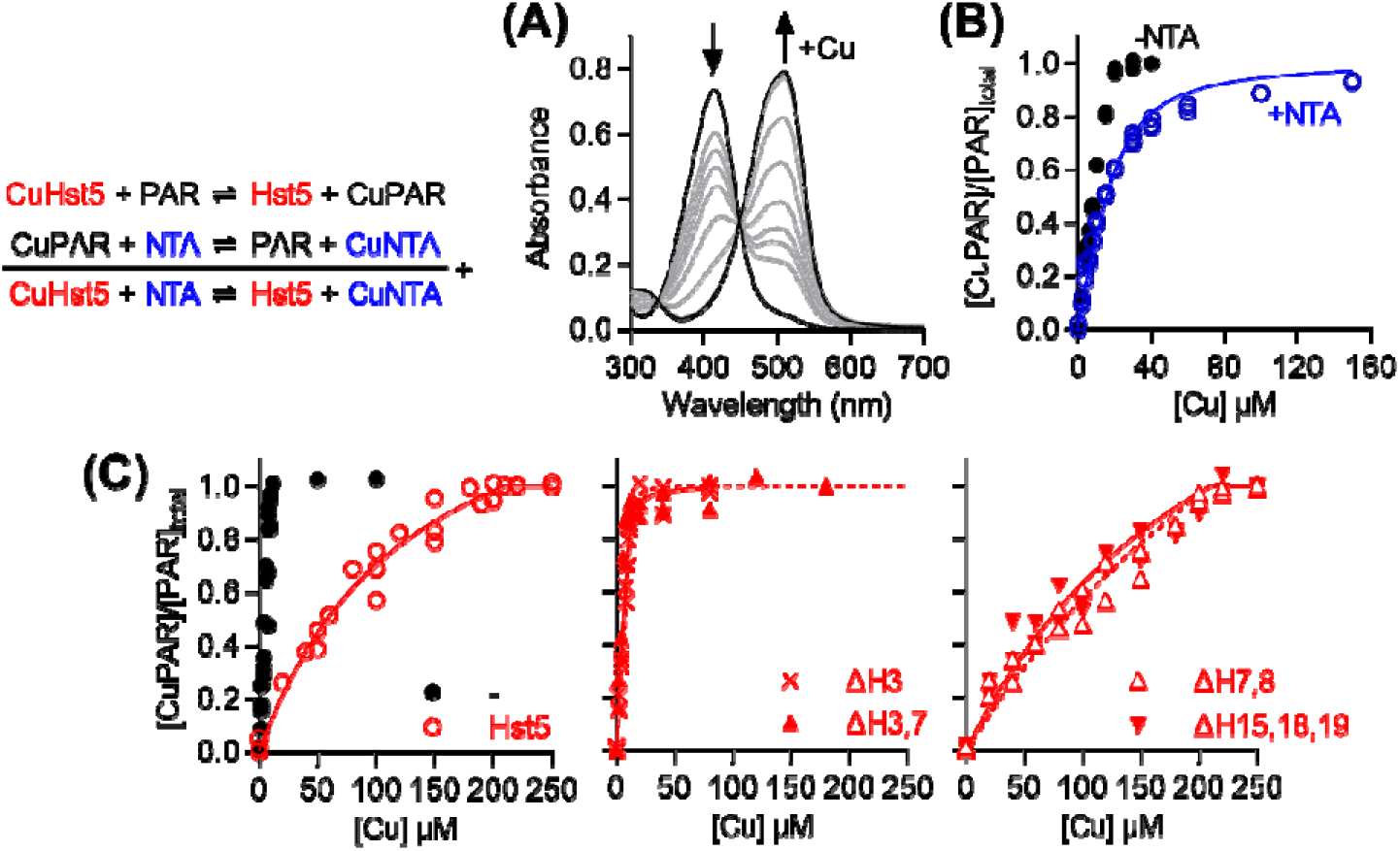
Cu(II) affinity of Hst5. **(A)** Representative spectral changes upon titration of Cu(II) (0–160 µM) into *apo*-PAR (20 µM). **(B)** Normalised plot of the absorbance intensities of CuPAR at 512 nm upon titration of Cu into *apo-*PAR (20 µM) in the absence (●) or presence (○) of NTA (400 µM). **(C)** Normalised plot of the absorbance intensities of CuPAR at 512 nm upon titration of Cu into *apo-*PAR (10 µM) in the absence or presence of Hst5 peptides (200 µM).

Consistent with an earlier report^17^, deletion of either *bis*-His motif did not affect the competition between Hst5 and PAR (Figure 10C), confirming that these residues do not participate in binding Cu(II). By contrast, deletion of His3 abolished the competition with PAR (Figure 10C), confirming the ATCUN motif as Cu(II)-binding ligands. More precise affinities for these variants were obtained *via* competition with the fluorometric Cu(II) chelator DP2^53^. By monitoring quenching of *apo*-DP2 fluorescence at 550 nm, log *K*_Cu_ values of 9.3 (±1.0) and 9.4 (±0.8) were obtained for the ΔH3 and ΔH3,H7 variants, respectively (Figure S9D). These values indicate that loss of the ATCUN His weakened the affinity of Hst5 to Cu(II) by ∼100-fold.

## DISCUSSION

### The role of metal binding in histatin activity: a framework for other metallo-AMPs

Our work establishes that Hst5 does not contribute to nutritional immunity against GAS, since this AMP does starve this bacterium of nutrient Zn, nor does it enhance Zn or Cu toxicity. These findings are consistent with the results from a genome-wide screen of a GAS mutant library, which did not identify Zn uptake, Zn efflux, or Cu efflux genes as essential for growth in saliva^54^.

The low affinity of Hst5 to Zn, particularly when compared with the high affinities of the Zn uptake lipoproteins AdcAI and AdcAII, explains why Hst5 does not starve GAS of *nutrient* Zn. Here, the antimicrobial protein calprotectin provides a useful comparison. Calprotectin binds two Zn ions with affinities (log *K*_Zn_ >11 and >9.6)^55^ that are comparable to those of AdcAI and tighter than that of AdcAII. Indeed, adding calprotectin induces a robust Zn starvation response in streptococci^56,57^, consistent with its established role in nutritional immunity.

Its low affinity to Zn also explains why Hst5 only weakly influences availability of *excess* (*toxic*) Zn to GAS. Like most culture media, our growth medium^20^ contains phosphate (∼6 mM) and amino acids (∼6 mM total), which would outcompete Hst5 (50 µM) for binding Zn^45^. For these reasons, synergistic effects between Zn and Hst5, such as those observed *in vitro* against *C. albicans*, may not result from a direct binding of Zn to Hst5. Instead, the separate biological effects of Zn and Hst5 may need to be considered. For instance, Zn and Hst5 may act on the same cellular targets or pathways. Alternatively, growth and survival of cells in the presence of Hst5 may require certain proteins that become poisoned in the presence of Zn (or *vice versa*).

If competing ligands become depleted, for example as a result of bacterial growth, then Hst5 can become competitive and bind Zn, particularly when Zn concentrations are high. Such shifts in Zn speciation likely explain why the protective effect of Hst5 on the GAS Δ*czcD* mutant during conditions of Zn stress became apparent only at the later stages of growth (*cf*. Figures 5F-G). The increased binding of Zn to Hst5 may at this point suppress non-specific Zn import into the GAS cytoplasm, for instance by outcompeting promiscuous divalent metal transporters or by suppressing direct Zn diffusion across the lipid bilayer.

Saliva contains ∼10 mM phosphate^58,59^ and proteinaceous components that may also bind Zn^60^. Unlike *in vitro* growth media, saliva and its components are continuously refreshed *in vivo*. Therefore, Hst5 is unlikely to strongly influence Zn speciation and availability in saliva. *In vivo*, synergistic effects between Zn and Hst5 may nonetheless occur, but likely *via* mechanisms that do not rely on formation of a Zn-Hst5 complex.

By contrast to Zn, the high affinity of Hst5 to Cu(II) explains why this AMP strongly influences Cu availability to GAS (*cf*. Figure 7) and, presumably, other microbes. Hst5 outcompetes background competing ligands that may also bind Cu, such as phosphate (log *K*_Cu_ ∼ 3.3^61^) and amino acids. In addition, Hst5 likely outcompetes transporters that catalyse *non-specific* Cu(II) uptake into GAS. This model will need to be tested by directly competing Hst5 and these transporters *in vitro*, but the latter are yet to be identified and are likely to be diverse.

Does Hst5 suppress *nutrient* Cu availability and cause Cu starvation? The GAS genome does not encode Cu-dependent proteins or enzymes, and so this bacterium is not thought to use or uptake nutrient Cu. Therefore, this proposal will need to be tested using other microbes that do need nutrient Cu. Nevertheless, parallels can again be drawn with calprotectin, which binds two Cu(II) ions with affinities (*log K*_Cu_ = 11.4 and 12.7)^62^ that are comparable to that of Hst5. Treatment with calprotectin induces a Cu starvation response in *C. albicans*^62^, suggesting that Hst5 may also elicit microbial Cu starvation response. Yet, Hst5 activity against this fungus appears linked to Cu excess and not Cu starvation^17^. Thus, the potential role of Hst5 in limiting nutrient Cu awaits further clarification.

The approaches described here can help define the role of metal binding in the function of metallo-AMPs in general. As an illustration, microplusin, a Cu-binding AMP from cattle ticks, is thought to withhold nutrient Cu from *Cryptococcus neoformans*^63,64^. This proposal was based on the observation that supplemental Cu suppressed the antimicrobial activity of this AMP. By measuring expression of Cu-responsive genes and total intracellular Cu levels in *C. neoformans*, one can determine whether microplusin binds Cu outside or inside target cells, and whether microplusin indeed influences Cu availability to these cells. By measuring the Cu affinity of microplusin and comparing it to those of key Cu uptake transporters in *C. neoformans* such as Ctr1, one can further determine whether microplusin is likely to bind Cu in the relevant host fluid, or whether the synergy between this AMP and Cu is associated with other unidentified mechanisms.

### Metal binding by histatins: implications for bacterial colonisation in the oral cavity and oropharynx

GAS causes >600 million worldwide cases of pharyngitis each year^65^, although asymptomatic carriage in the oropharynx is common, especially among children^66^. This host niche is rich in saliva, and the interactions between GAS and components of this host fluid are key for colonisation, infection, and subsequent transmission of this bacterium^67-69^. For example, exposure to saliva promotes aggregation of GAS and blocks adherence to mucosal epithelia^70^. However, GAS produces surface adhesins that aid in binding to host mucosal surfaces^71^. Saliva also contains polysaccharides and glycoproteins that may serve as sources of nutrients. Accordingly, carbohydrate utilisation genes in GAS are highly expressed upon exposure to saliva^72^, and mutant strains lacking these genes show decreased fitness in saliva^54^. Given the widely reported antimicrobial activity of Hst5, salivary histatins are thought to act as antimicrobial peptides. Yet, our work shows that Hst5 does not kill or inhibit growth of GAS under saliva-relevant conditions.

Discussions surrounding histatins have thus far focused on the *antagonistic* relationship between these AMPs and opportunistic oral pathogens *in vivo*. By comparison, little is known about the potentially *harmonious* relationship between histatins and microbes that normally colonise healthy oral and oropharyngeal tissues. We showed here that Hst5 does not kill oral streptococci, which represent the most abundant microbial taxon in healthy human oral cavity^73-76^ and oropharynx^77^. This lack of an anti-streptococcal effect contrasts with the potent antibacterial effects of Hst5 against ESKAPE pathogens^11^, although the latter are worth revisiting, to verify that they are not associated with artificial experimental conditions that do not mimic the saliva.

While total levels of intact, full-length Hst5 and major histatins in fresh salivary gland secretions are high (up to 50 μM)^9^, steady-state levels in whole saliva are low^9^ as a consequence of peptide degradation by unidentified salivary proteases and proteases from resident oral microbes^78,79^. Nearly fifty histatin-derived peptide fragments have been identified^80,81^. Many are associated with reduced antimicrobial activities^8,80^, raising the question whether an antimicrobial role is the major physiological role for the histatins.

Intriguingly, proteolytic cleavage of histatins in saliva typically leads to retention of the original Cu(II)-binding ATCUN motif (DSH-) and simultaneous generation of new ATCUN motifs as byproducts (RHH-, EKH-, KFH-, KRH-, KHH-, HSH-)^80^. These diverse new motifs likely continue to bind Cu(II)^82^, raising the intriguing possibility that Cu binding is the key physiological role for histatins.

We speculate that histatins contribute to oral and oropharyngeal health by buffering Cu availability. Steady-state levels of Cu in healthy saliva are sub-stoichiometric relative to histatins^41-43^, but additional Cu does enter the oral cavity through food (*e*.*g*. liver, shellfish, dark chocolate). In addition, Cu levels in saliva can also rise during periodontal diseases^83-85^. By buffering Cu, histatins may protect resident oral microbes from the potential toxic effects of a sudden or sustained exposure to *excess* Cu, and thus promote microbial homeostasis in saliva-rich host niches.

Streptococci do not use nutrient Cu, and so these bacteria will only benefit from the action of histatins as Cu-buffering agents. This idea is not inconsistent with the relative dominance of *Streptococcus* species in the human oral cavity^74,75^ and oropharynx^77^. However, other resident oral microbes, such as commensal *Neisseria* species and even *C. albicans*, need nutrient Cu for respiration and energy production. Do histatins buffer *nutrient* Cu availability to these microbes? The oral cavity and oropharynx are also major entry points for pathogens that can cause oral, gut, and respiratory infections, many of which also need nutrient Cu. How is Cu availability managed, such that toxicity is limited to resident microbes but enhanced to foreign, potentially pathogenic microbes, and that nutrient supply is maintained to resident microbes but suppressed to pathogenic ones? Are there species-specific differences? What is the molecular basis of such differences? These studies are ongoing in our laboratory.

## METHODS

### Data presentation

Except growth curves, individual replicates from microbiological experiments are plotted, with shaded columns representing the means and error bars representing standard deviations. Growth curves show the means, with shaded regions representing standard deviations. The number of biological replicates (independent experiments using different starter cultures and performed on different days; *N*) is stated in figure legends. Statistical analyses have been performed on all data but notations of statistical significance are displayed on graphical plots only if they aid in rapid, visual interpretation. Unless otherwise stated, statistical tests used two-way analysis of variance using the statistical package in GraphPad Prism 8.0. All analyses were corrected for multiple comparisons. In the case of metal-protein and metal-peptide titrations, individual data points from two technical replicates (independent experiments performed on different days but using the same protein or peptide preparation) are plotted, but only representative spectra are shown for clarity of presentation.

### Reagents

Sulfate and chloride salts of metals were used interchangeably. Peptides were synthesised commercially as the acetate salt, purified to >95% (GenScript), and confirmed to be metal-free by ICP MS. Concentrations of stock peptide solutions were estimated using solution absorbances at 280 nm in Mops buffer (50 mM, pH 7.4; ε_280_ = 2667 cm^-1^). Concentrations of fluorometric and colourimetric metal indicators (Zincon, PAR, Magfura-2, Quin-2, BCS, DP-2) were standardised using a commercial standard solution of copper chloride. Concentrations of optically silent chelators (NTA) were standardised by competition with a standardised solution of Zn-Zincon.

### Strains and culture conditions

Bacterial strains are listed in Dataset S1b. All bacterial strains (Dataset S1b) were propagated from frozen glycerol stocks onto solid THY (Todd Hewitt + 0.2% yeast extract) medium without any antibiotics. Liquid cultures were prepared in THY or CDM-glucose^20^. All solid and liquid growth media contained catalase (50□μg/ml).

### Survival assays

Fresh colonies from an overnight THY agar were resuspended to 10^6^-10^7^ CFU/mL in either potassium phosphate buffer (10 mM, pH 7.4), Mops buffer (10 mM, pH 7.4), or artificial salivary salts (pH 7.2; Dataset S1a). The cultures were incubated at 37 °C with or without Hst5 and/or metals as required. At *t* = 0, 1, and 3 h, cultures were sampled and serially diluted in CDM-glucose. Exactly 10 µL of each serial dilution was spotted onto fresh THY agar. Colonies were enumerated after overnight incubation at 37 °C.

### Growth assays

Colonies from an overnight THY agar were resuspended in CDM-glucose to an OD_600_ = 0.01. Growth was assessed at 37 °C in flat-bottomed 96-well plates (200 µL per well) using an automated microplate shaker and reader. Each plate was sealed with a gas permeable, optically clear membrane (Diversified Biotech). OD_600_ values were measured every 20□min for 10 h. The plates were shaken immediately before each reading (200□rpm, 1□min, double orbital mode). OD_600_ values were not corrected for path length (*ca*. 0.58□cm for a 200-μl culture).

### RNA extraction

Colonies from an overnight THY agar were resuspended in CDM-glucose to an OD_600_ = 0.01 and incubated in 24-well plates (1.6 mL per well) without shaking at 37 °C. Each plate was sealed with a gas permeable, optically clear membrane (Diversified Biotech). At *t* = 4 h, cultures were centrifuged (4,000 x *g*, 4°C, 5 min) and bacterial pellets were resuspended immediately in RNAPro Solution (0.5 mL; MP Biomedicals). Bacteria were lysed in Lysing Matrix B and total RNA was extracted following manufacturer’s protocol (MP Biomedicals). Crude RNA extracts were treated with RNase-Free DNase I (New England Biolabs). Complete removal of gDNA was confirmed by PCR using gapA-check-F/R primers (Dataset S1c). gDNA-free RNA was purified using Monarch RNA Clean-up Kit (New England Biolabs) and visualised on an agarose gel.

### qRT-PCR analyses

cDNA was generated from RNA (1.6 µg) using SuperScript® IV First-Strand Synthesis System (Invitrogen). Each qRT-PCR reaction (20 µL) contained cDNA (5 ng) as template and the appropriate primer pairs (0.4 µM; Dataset S1c). Samples were analysed in technical duplicates. Amplicons were detected with Luna® Universal qRT-PCR Master Mix (New England Biolabs) in a CFXConnect Real-Time PCR Instrument (Bio-Rad Laboratories). *C*_q_ values were calculated using LinRegPCR^86^ after correcting for amplicon efficiency. *C*_q_ values of technical duplicates were typically within ± 0.25 of each other. *holB*, which encodes DNA polymerase III, was used as reference gene. Its transcription levels remained constant in all of the experimental conditions tested here.

### Intracellular metal content

Colonies from an overnight THY agar were resuspended in CDM-glucose to an OD_600_ = 0.02 and incubated at 37 °C with or without Hst5 and/or metals as required. At *t* = 4 h, an aliquot was collected for the measurement of plating efficiency (colony counts). The remaining cultures were harvested (5,000 g, 4 °C, 10 min), and washed once with ice-cold wash buffer (1 M *D*-sorbitol, 50 mM Tris-HCl, 10 mM MgCl_2_, 1 mM EDTA, pH 7.4) and twice with ice-cold PBS. The final pellets were dissolved in concentrated nitric acid (100 µL), heated (85 °C, 1.5 h), and diluted to 3.5 mL with 2 % nitric acid. Total metal levels were determined by ICP MS. The results were normalised to colony counts.

### Elution of Zn-Hst5 on a desalting column

*Apo*-Hst5 (100 µM) was incubated with 1.5 molar equiv. of Zn for 15 min at the bench and loaded onto a polyacrylamide desalting column (1.8 kDa molecular weight cutoff, Thermo Scientific). Peptide content in each fraction was verified using QuantiPro BCA Assay Kit (Merck). Zn content was determined using PAR against a standard curve.

### Equilibrium competition reactions

Our approach to determine metal-binding affinities followed that described by Young and Xiao^47^. For each competition (eq 1 below), a master stock was prepared to contain both competing ligands (L1 and L2) in Mops buffer (50 mM, pH 7.4). Serial dilutions of the metal (M) were prepared separately in deionised water. Exactly 135 µL of the master stock was dispensed into an Eppendorf UVette and 15 µL of the appropriate metal stock was added. Solution absorbances were used to calculate concentrations of *apo*- and metalated forms of the relevant ligand. These concentrations were plotted against metal concentrations and fitted in DynaFit^87^ using binding models as described in the text. The known association or dissociation constants for all competitor ligands are listed in Dataset S1d:

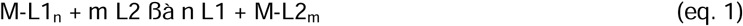

### Overexpression and purification of AdcAI and AdcAII

Nucleic acid sequences encoding the soluble domains of AdcAI (from Thr21) and AdcAII (from Thr31) from M1GAS strain 5448 were subcloned into vector pSAT1-LIC using primers listed in Dataset S1c. This vector generates N-terminal His6-SUMO fusions with the target ORF. The resulting plasmids were propagated in *E. coli* Dh5α, confirmed by Sanger sequencing, and transformed into *E. coli* BL21 Rosetta 2(DE3).

To express the proteins, transformants were plated onto LB agar. Fresh colonies were used to inoculate LB (1 L in 2 L baffled flasks) to an OD_600_ of 0.01. The culture media contained ampicillin (100 µg/mL) and chloramphenicol (33 µg/mL) as required. Cultures were shaken (200 rpm, 37 °C) until an OD_600_ of 0.6–0.8 was reached, and expression was induced by adding IPTG (0.1 mM). After shaking for a further 16 h at 20 °C, the cultures were centrifuged (4000 × *g*, 4 °C) and the pellets were resuspended in buffer A500 (20 mM Tris–HCl, pH 7.9, 500 mM NaCl, 5 mM imidazole, 10% glycerol).

To purify proteins, bacteria were lysed by sonication (40 kpsi), centrifuged (20,000 × *g*, 4°C), and filtered through a 0.46 µm PES membrane filtration unit. Clarified lysates were loaded onto a HisTrap HP column (Cytiva). The column was washed with 10 column volumes (CV) of buffer A500 followed by 10 CV of buffer A100 (20 mM Tris–HCl, pH 7.9, 100 mM NaCl, 10% w/v glycerol) containing imidazole (5 mM). Both AdcAI and AdcAII were bound to the column and subsequently eluted with 3 CV of buffer A100 containing 250 mM imidazole followed by 5 CV of 500 mM imidazole.

Protein-containing fractions were loaded onto a Q HP column (Cytiva). The column was washed with 5 CV of buffer A100 and bound proteins were eluted using a step gradient of 0, 10, 15, and 20% buffer C1000 (20 mM Tris–HCl, pH 7.9, 1000 mM NaCl, 10% w/v glycerol). Eluted proteins were incubated overnight at 4 °C with hSENP2 SUMO protease to cleave the His6-SUMO tag from the target protein. Samples were passed through a second Q HP column and the flowthrough fractions containing untagged target protein were collected.

## AUTHOR CONTRIBUTIONS

KD conceived the project with input from NJ, SC. JH, KD, LS, NJ designed experiments. NJ provided oral streptococci strains. IH, SC, YS synthesised peptides for preliminary studies. IH, JH, KD, LS examined the effects of peptides on bacterial growth. JB, KD measured gene expression by qRT-PCR. KD, LS measured metal levels by ICP MS. IH, JH, SF measured affinities of peptides to Cu and Zn. JH produced AdcA and AdcAII proteins, and measured their affinities to Zn. LS examined the effects of peptides on bacterial survival. JH, KD, LS prepared figures and drafted the manuscript.

## ACKNOWLEDGEMENT

This work was funded by the Wellcome Trust grant number 214930/Z/18/Z. For the purpose of open access, the author has applied a CC BY public copyright licence to any Author Accepted Manuscript version arising from this submission. This project was also supported by a Flexible Funding Award from the Durham Biophysical Sciences Institute. SF and JB were supported by studentships from the BBSRC Newcastle-Liverpool-Durham Doctoral Training Partnership.

T Blower (Durham University) provided constructs and reagents for production of AdcAI and AdcAII. GAS Δ*adcAI* and Δ*czcD* mutant strains were from C Ong, A McEwan, and M Walker (The University of Queensland). Quin2 was from T Young (Durham University). We thank R Borthwick and P Chivers (Durham University), and Y Ahmed and J Quinn (Newcastle University Biosciences Institute) for insightful discussions.

