## Supporting Figures for "The role of metal binding in the function of the human salivary antimicrobial peptide histatin-5"

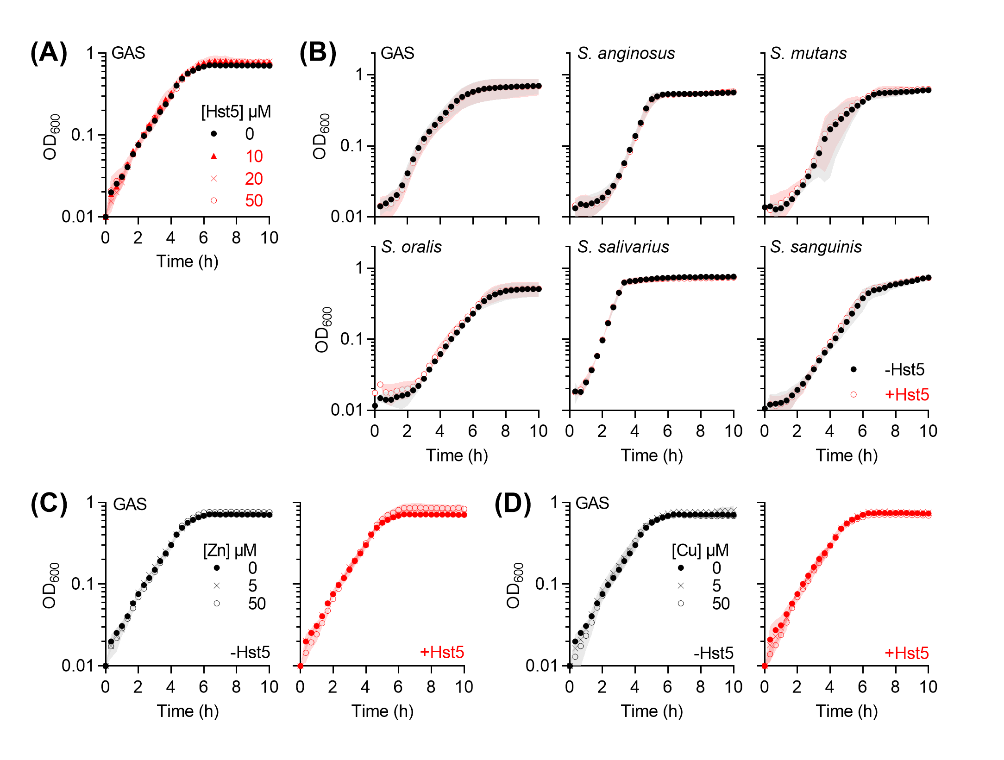

**Figure S1. Effects of Hst5 in THY.**

**(A) Direct effects against GAS.** Bacteria (*N =* 2) were cultured in the presence of Hst5 (0, 10, 20, 50 µM).

**(B) Direct effects against oral streptococci.** Bacteria (*N =* 2) were cultured with or without Hst5 (50 µM).

**(C) Zn-dependent effects against GAS.** Bacteria (*N =* 2) were cultured in the presence of Zn (0, 5, 50 µM), with or without Hst5 (50 µM).

**(D) Cu-dependent effects against GAS.** Bacteria (*N =* 2) were cultured in the presence of Cu (0, 5, 50 µM), with or without Hst5 (50 µM).

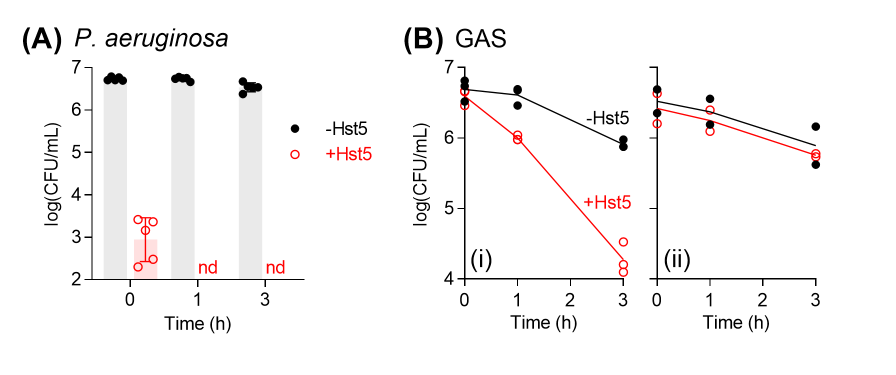

**Figure S2. Bactericidal effects of Hst5.**

**(A) Bactericidal effects of Hst5 against *P. aeruginosa*.** Bacteria (~5 x 10^6^ CFU/ml, *N =* 5) were incubated in potassium phosphate buffer (10 mM, pH 7.4) with or without Hst5 (50 µM). nd, not detected (detection limit log(CFU/ml) = 2). Note that at *t* = 0 h, approximately 5 min passed between addition of Hst5 into bacterial cultures and plating out for enumeration.

**(B) Carryover salts in the inoculum abolished the bactericidal effects of Hst5 against GAS in Mops.** Bacteria (~5 x 10^6^ CFU/ml, *N =* 3) were incubated in Mops (10 mM, pH 7.4) with or without Hst5 (50 µM). The inoculum were prepared in: **(i)** Mops (10 mM, pH 7.4) or **(ii)** potassium phosphate buffer (10 mM, pH 7.4).

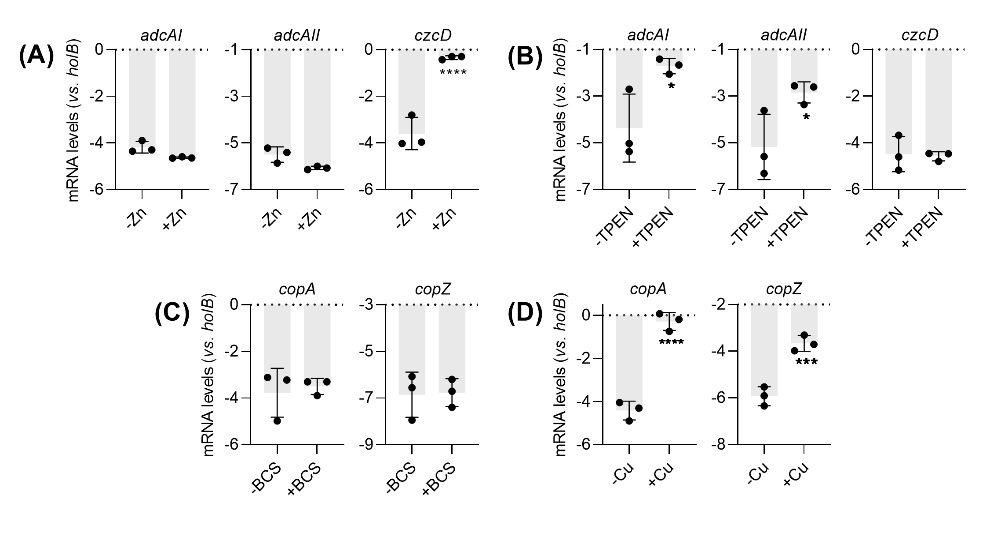

**Figure S3. Effects of (A) Zn, (B) TPEN, (C) BCS, and (D) Cu on gene expression in wild-type GAS.**

**(A)-(B)** Bacteria (*N =* 3) were cultured in CDM-glucose for 4 h with or without added Zn (5 µM) or TPEN (100 nM) as indicated. Levels of *adcAI*, *adcAII*, and *czcD* mRNA were determined by qRT-PCR and normalised to *holB*. Growth in the presence of Zn led to upregulation of *czcD* (*****P* < 0.0001) but not *adcAI* (*P* = 0.3) or *adcAII* (*P* = 0.2). Growth in the presence of TPEN led to upregulation of *adcAI* (**P* = 0.01) and *adcAII* (**P* = 0.03) but not *czcD* (*P* = 1.0).

**(C)-(D)** Bacteria (*N =* 3) were cultured in CDM-glucose for 4 h with or without added Cu (1 µM) or BCS (250 µM) as indicated. Levels of *copA* and *copZ* mRNA were determined by qRT-PCR and normalised to *holB*. Growth in the presence of BCS did not perturb expression of *copA* (*P* = 0.9) and *copZ* (*P* = 1.0). Growth in the presence of Cu led to upregulation of *copA* (*****P* < 0.0001) and *copZ* (****P* = 0.0002)*.*

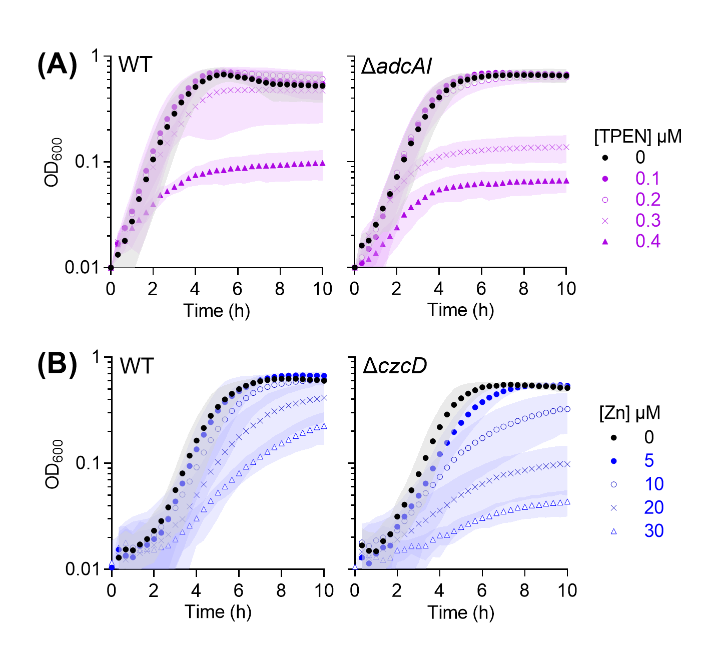

**Figure S4. Characteristic phenotypes of the ∆*adcAI* and ∆*czcD* mutant strains.**

**(A) TPEN-sensitive phenotype of the ∆*adcAI* mutant.** Bacteria (*n* = 3) were cultured in CDM-glucose in the presence of TPEN (0 – 0.4 µM).

**(B)** **Zn-sensitive phenotype of the ∆*czcD* mutant.** Bacteria (*N =* 3) were cultured in CDM-glucose in the presence of added Zn (0 – 30 µM).

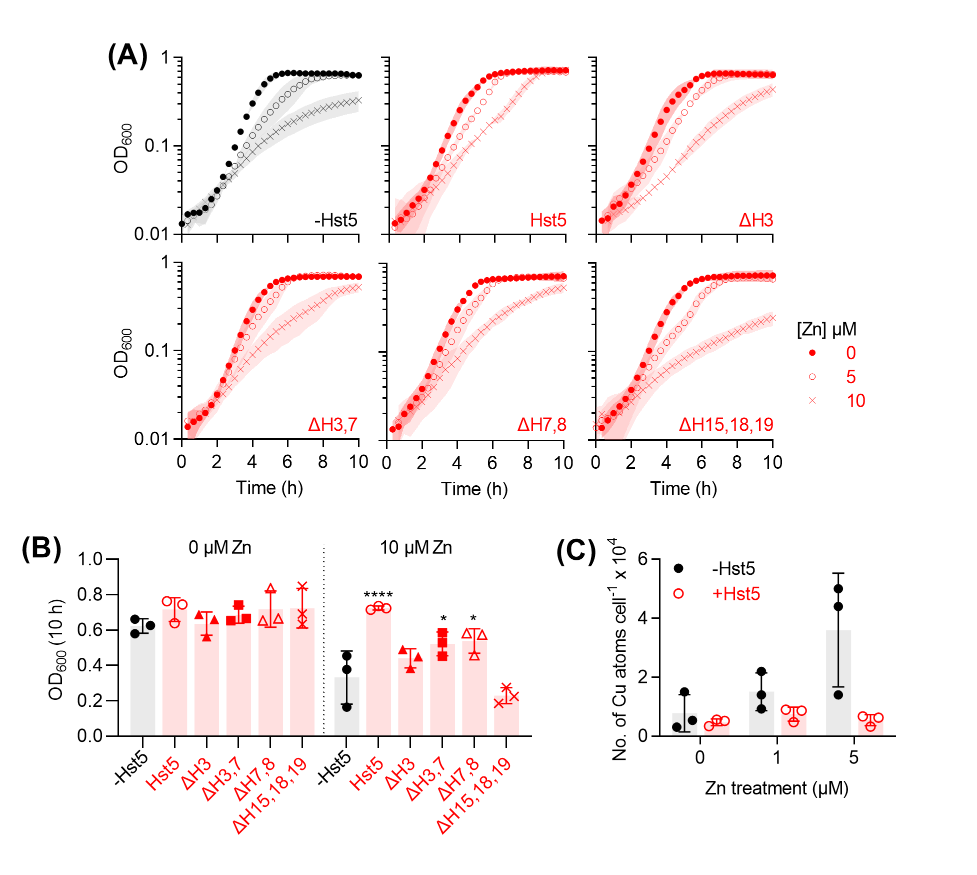

**Figure S5. Effects of Hst5 on the ∆*czcD* mutant.**

**(A)-(B)** **Effects of Hst5 variants on growth. (A)** GAS ∆*czcD* mutant strain (*N =* 3) was cultured in CDM-glucose in the presence of added Zn (0, 5, 10 µM), with or without Hst5 or its variants (50 µM each; *cf.* Table 1 for peptide sequences). **(B)** Plot of OD_600_ values at *t* = 10 h from growth curves in panel A. The following peptides increased final culture densities: Hst5 (*****P* < 0.0001), ∆H3,7 (**P* = 0.04), and ∆H7,8 (**P* = 0.03). The following peptides had no effect: ∆H3 (*P* = 1.0), ∆H15,18,19 (*P =* 0.4).

**(C) Effects of Hst5 on intracellular Cu levels.** GAS ∆*czcD* mutant strain (*N =* 3) was cultured in CDM-glucose in the presence of added Zn (0, 1, 5 µM), with or without Hst5 (50 µM). Total intracellular Cu levels were measured by ICP MS analyses.

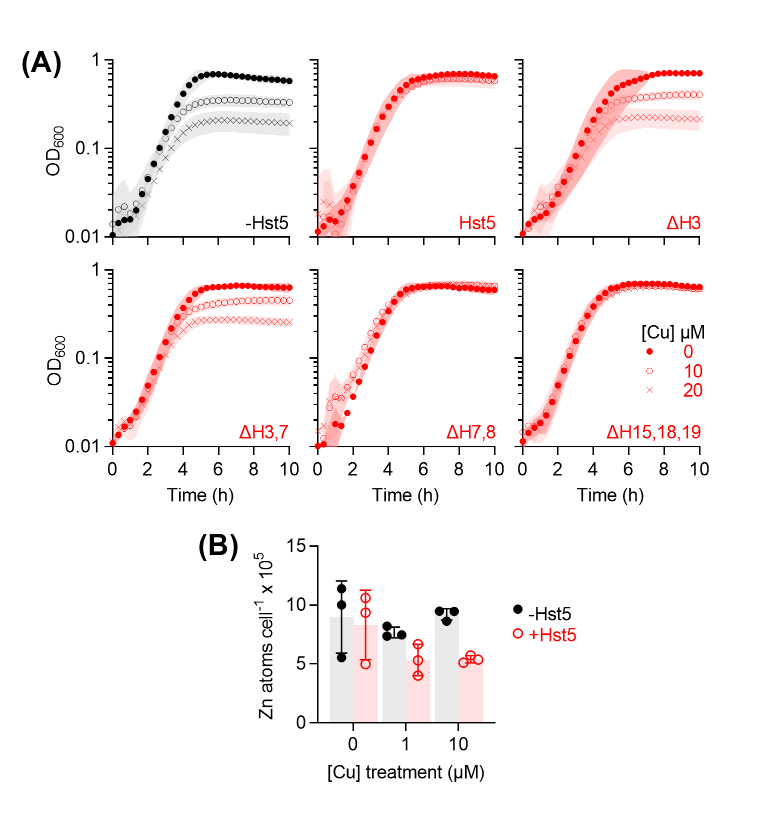

**Figure S6. Effects of Hst5 on the ∆*copA* mutant.**

**(A) Effects of Hst5 variants on growth.** GAS ∆*copA* mutant strain (*N =* 3) was cultured in CDM-glucose in the presence of added Cu (0, 10, 20 µM), with or without Hst5 or its variants (50 µM each; *cf.* Table 1 for peptide sequences).

**(B) Effects of Hst5 on intracellular Zn levels.** GAS ∆*copA* mutant strain (*N =* 3) was cultured in CDM-glucose in the presence of added Cu (0, 1, 10 µM), with or without Hst5 (50 µM). Total intracellular Zn levels were measured by ICP MS analyses.

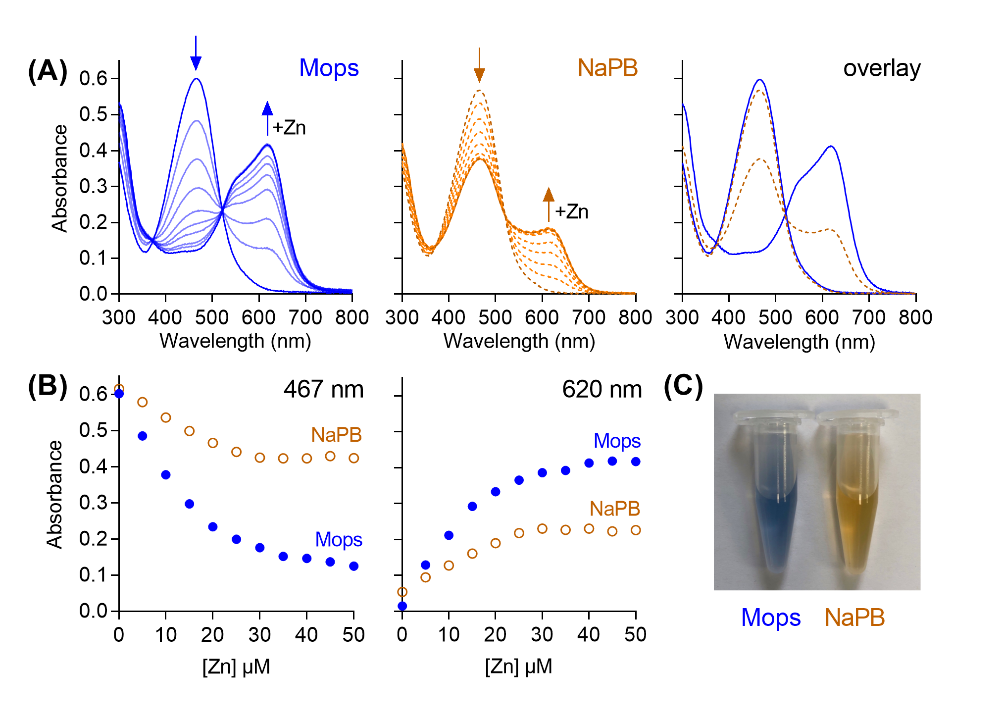

**Figure S7. Phosphate buffer competes with Zincon for Zn.**

**(A)** Representative spectral changes upon titration of Zn (0 – 50 µM) into *apo*-Zincon (20 µM) in Mops buffer (50 mM, pH 7.4; solid traces) or sodium phosphate buffer (50 mM, pH 7.4; NaPB, dashed traces).

**(B)** Plot of absorbance values at 467 nm or 620 nm from panel A.

**(C)** Loss of the characteristic blue colour of Zn-Zincon complex upon incubation in NaPB for >10 min.

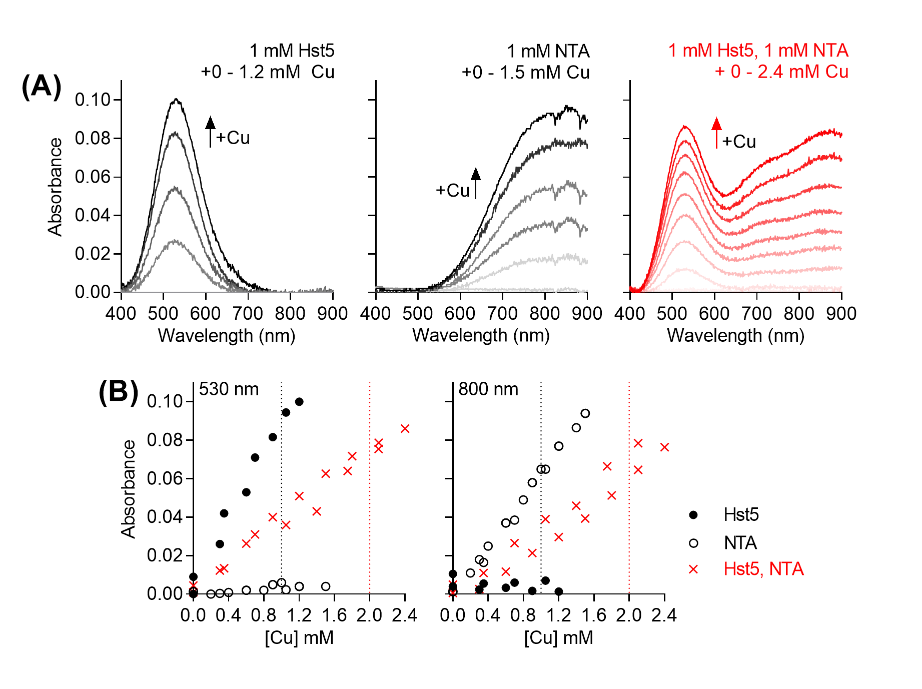

**Figure S8. Competition between Hst5 and NTA for Cu(II).**

**(A)** Representative spectral changes upon titration of Cu into Hst5 (1 mM, left panel), NTA (1 mM, middle panel), or a mixture of Hst5 and NTA (1 mM each, right panel) in Mops buffer (50 mM, pH 7.4). Upward arrows indicate increases in absorbance intensity.

**(B)** Plot of absorbance values from panel A, demonstrating the lack of end-points at 530 nm and 800 nm, which report the binding of Cu into Hst5 and NTA, respectively. Vertical dotted lines indicate the Cu concentrations where end-points are expected.

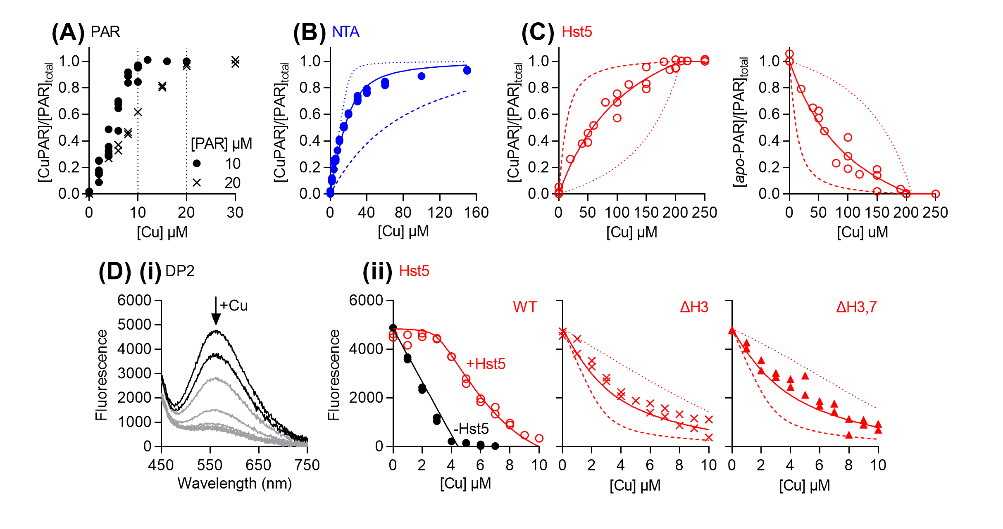

**Figure S9. Cu(II) affinities of Hst5 and its variants.**

**(A) Direct titration of Cu(II) into PAR.** Two PAR concentrations were used. Both titrations confirmed saturation of PAR at 1 molar equiv. of Cu(II).

**(B) Competition between PAR and NTA.** Competition curves showing the formation of CuPAR upon addition of Cu (0 – 160 µM) into a mixture of PAR (20 µM) and NTA (400 µM). The curve fit (solid line) yielded log *K*_Cu_ = 12.8 for PAR, which clearly departs from simulations of 10X higher (dotted line) and lower (dashed line) affinity.

**(C) Competition between PAR and Hst5.** Competition curves showing the formation of CuPAR (left panel) or the disappearance of *apo*-PAR (right panel) upon addition of Cu (0 – 250 µM) into a mixture of PAR (10 µM) and Hst5 (200 µM). Curve fits (solid lines) yielded log *K*_Cu_ = 12.1 for Hst5, which clearly depart from simulations of 10X higher (dotted lines) and lower (dashed lines) affinities.

**(C) Competition between DP2 and Hst5. (i)** Changes in the fluorescence spectra upon addition of Cu (0 – 8 µM) into a solution of *apo*-DP2 (4 µM). **(ii)** Competition curves between DP2 (4 µM) and Hst5 (3 µM), the ∆H3 variant (8 µM), or the ∆H3,7 variant (8 µM). The data confirmed that Hst5 outcompeted DP2. Competition curves with the Hst5 variants were best fitted to a 2-site model where both Cu atoms bind with equal affinities, suggesting the presence of a second, weak binding site. Curve fits (solid lines) yielded log *K*_Cu_ = 9.3 and 9.4 for the ∆H3 and ∆H3,7 variants, respectively, which clearly depart from simulations of 10X higher (dotted lines) and lower (dashed lines) affinities.
